# Unraveling the Complexity of Abdominal Aortic Aneurysm: Multiplexed Imaging Insights into C-Reactive Protein-Related Variations

**DOI:** 10.1101/2024.02.22.581315

**Authors:** Eun Na Kim, Hee Young Seok, Jiwon Koh, Wookyeom Yang, Gyu Ho Lee, Woo Hee Choi, Joon Seo Lim, You Jung Ok, Jae-Sung Choi, Chong Jai Kim, Lizhe Zhuang, Young Hwan Chang, Se Jin Oh

## Abstract

**Background:** Abdominal aortic aneurysm (AAA) is a potentially lethal condition that often remains asymptomatic until it ruptures. Recent research suggests that immune-inflammatory processes are associated with AAA development, yet the exact mechanisms remain unclear. Serum C-reactive protein (CRP) serves as a prognostic marker for AAA and various cardiovascular diseases. When CRP accumulates in damaged tissues, it transforms into a monomeric form, exacerbating tissue damage. Our previous study confirmed the presence of CRP deposition in eroded AAA and atherosclerosis regions, accompanied by an increased infiltration of inflammatory cells. However, a comprehensive understanding of the specific changes in the inflammatory cellular landscape attributable to CRP deposition is lacking. Here, we aimed to explore cellular-level alterations in AAAs associated with varying CRP levels.

**Methods:** We categorized AAA patients into High-CRP (≥0.1 mg/dL, and ≥3+ CRP IHC score, n=6) and Low-CRP (≤0.1 mg/dL, and ≤1+ CRP IHC score, n=3), and used normal aorta specimens as a baseline control. The cellular landscape of immune and stromal components was characterized using Co-Detection by Indexing (CODEX) tissue imaging with 31 DNA-barcoded antibodies, followed by single-cell-based analysis and GPU-accelerated unsupervised clustering.

**Results:** We identified 51 distinct immune and stromal cell types in the cohort and revealed significant differences in protein expression patterns among the groups. In AAA, stromal cells decreased significantly, while immune cell proportions sharply increased. Lag3+ T cell regulators decreased, leading to an increase in CD3+ T cells. The composition of immune cells within atherosclerotic plaques was associated with the degree of CRP deposition in AAA. The High-CRP group showed increased M1 and Ki67+ proliferating macrophages, while the Low-CRP group exhibited intensified fibrosis with M2 macrophages.

**Conclusions:** Our study found significant variations in immune cell distribution within AAA walls based on CRP levels. These findings suggest a potential link between CRP-related immune changes and AAA progression.

**HIGHLIGHTS:**

- By performing CODEX (co-detection by indexing) multiplexed imaging on paraffin-embedded, formalin-fixed archived abdominal aortic aneurysm (AAA) tissue samples with varying CRP levels, we segmented 415,365 cells into 51 distinct clusters.
- In AAA, the presence of CRP deposition led to variations in the spatial relationships, distances, and enrichment patterns among immune and stromal cells..
- In AAA-High CRP, there was an intensified infiltration of M1 macrophages and various immune cells observed in the atherosclerotic plaque.
- In AAA-Low CRP, severe stromal fibrosis was associated with M2 macrophages.
- Our results suggest that the deposition of CRP into the atherosclerotic plaque alters the immune-stromal landscape in AAA.

## INTRODUCTION

Abdominal aortic aneurysm (AAA) is a life-threatening condition characterized by the gradual expansion of the abdominal aorta, often without noticeable symptoms until the point of imminent rupture. Sampson et al. reported that AAA is often detected after it has ruptured, resulting in a high fatality rate of 60% upon rupture, emphasizing its critical nature and the need for early detection and intervention^1^. Despite the severity of the condition, many aspects of the pathogenesis of AAA development and progression remain poorly understood. Furthermore, effective pharmacological interventions to halt the progression of AAA have not yet been developed. A comprehensive understanding of the mechanisms driving AAA deterioration is urgently needed to improve the current situation.

In recent years, there has been an increasing recognition of the role of autoimmune mechanisms and immune-inflammatory pathways in the onset and progression of AAA^2^. The critical cell populations involved in AAA pathogenesis include B and T lymphocytes, macrophages, mast cells, and natural killer cells^3^. Notably, an upregulation of Th1 mononuclear responses has been observed, leading to increased secretion of cytokines such as interleukin-2, interferon-gamma, and tumor necrosis factor-alpha from macrophages. This proinflammatory cytokine profile promotes the secretion of proinflammatory osteopontin in vascular smooth muscle cells, thereby exacerbating the inflammatory response in the vessel wall^4–6^. The concentration of CD4+ T cells and interleukin-17 significantly increases in AAA tissues within the periadventitial vascular-associated lymphatic tissue, as reported by Chang *et al.*^7^ These findings emphasize the crucial role that immune and inflammatory mechanisms play in the development of AAA.

Furthermore, increases in serum C-reactive protein (CRP) levels are evident in various cardiovascular diseases and are consistently associated with unfavorable prognoses. While the pentameric form of serum CRP in the bloodstream plays an anti-inflammatory and protective role, it undergoes a structural transformation into a monomeric form upon encountering damaged tissue membranes^8^. This transition leads to the deposition of CRP within the damaged tissue, where it assumes a pro-inflammatory role, ultimately accelerating tissue damage^9^.

In our previous study, we reported the deposition of CRP at the interface between the eroded aneurysmal vascular wall and atheromatous plaque, along with inflammatory cell infiltration in the atheroma of AAA, a phenomenon not typically observed in aortic dissection or the normal aorta^10^. In AAA specimens with high serum CRP levels, CRP deposition was more prominent and associated with a higher density of inflammatory cells, including CD68+ macrophages, as well as complement deposition, compared to AAA specimens with lower serum CRP levels, which had weak and focal CRP deposition at the aneurysmal wall. Additionally, we found that the presence of CRP deposition in the atheroma of AAA leads to a significant modification of the proteomic profile related to acute phase response signaling, atherosclerosis signaling, prothrombin activation pathway, and complement system within the aneurysmal tissue^6^. This suggests a potential association between CRP deposition and the progression of AAA. However, the phenotype of immune cells within the AAA vessel wall and the resulting composition of immune cells when CRP deposits within AAA atheroma remain poorly understood. There is a need to understand the complex interactions between CRP deposition, immune cells, and the microenvironment of vascular tissue in the context of AAA.

The intricate interplay between cancer cells and the surrounding microenvironment, especially interactions involving immune and stromal cells, is increasingly recognized as a pivotal factor in understanding disease prognosis and therapy^11^. The rapid development of multiplexed tissue imaging (MTI) techniques^12^ enables comprehensive and quantitative assessment of the microenvironmental landscape at the single-cell level. These methodologies facilitate the concurrent staining of diverse markers, precise quantification of specific cell phenotypes, and analysis of their spatial distribution, offering valuable insights at the single-cell level.

In this study, we utilized the CODEX (CoDetection by indexing)^13^ multiplexed tissue imaging technique to analyze the high-dimensional phenotypes of immune and stromal cells within AAA at a single-cell level. Our investigation focused on understanding how the deposition of CRP influences the multicellular spatial organization within AAA and alters the microenvironment of aortic aneurysms, with a particular focus on the interactions between inflammatory cells and stromal elements. Notably, we identified distinct variations in the composition of atheromatous plaques, aortic smooth muscle cells, and the spatial organization of immune cells within AAA associated with CRP deposition. These findings provide valuable insights into the dynamic interplay between CRP, the distribution of immune cells, and the microenvironment of abdominal aortic aneurysms, shedding light on this complex vascular pathology.

## METHODS

### Aorta Specimens

We used archived samples of AAA from patients who had undergone surgery at two tertiary referral centers in Seoul, South Korea (Table S1). Patients with systemic conditions that can result in elevated serum CRP levels and potentially worsen aortic inflammation, such as IgG4-related disease, Takayasu arteritis, and giant cell arteritis, were excluded from the study. Serum CRP levels were assessed just before surgery, and immunohistochemistry (IHC) staining was performed on the surgically resected aortic tissue using anti-CRP and anti-mCRP antibodies. The CRP IHC scoring method was conducted following the protocol established in our previous study on aortic aneurysms^10^. Based on the serum CRP level and the intensity of CRP immunostaining in the aneurysmal tissue, we classified patients into two groups: the High-CRP group (patient n = 6, tissue core n = 12) with serum CRP levels ≥ 0.1 mg/dL and ≥ 3+ CRP IHC score, and the Low-CRP group (patient n = 3, tissue core n = 5) with serum CRP ≤ 0.1 mg/dL and ≤ 1+ CRP IHC score. As a control group, we used healthy aorta specimens obtained during heart transplantation (patient n = 3, tissue core n = 6). Representative images of anti-CRP antibody IHC staining for the Low-CRP and High-CRP groups are shown in Figure S1.

### Tissue Microarray Construction

We created tissue microarrays (TMAs) from surgically resected aortic specimens to streamline analysis. A cardiovascular pathologist (ENK) annotated regions of interest, including atheroma, thinned aortic walls, and atheroma-eroded aortic intima, on the corresponding H&E slides. These annotations guided the construction of TMAs.

Cores measuring 3 mm in diameter were extracted from the identified regions and processed as TMAs. Specifically, areas encompassing atheromatous plaques and the interface between the aortic walls were chosen for core punching; specifically, 12 cores were taken from the High-CRP group, five cores from the Low-CRP group, and 6 cores from the normal aorta. The tissue microarrays were sectioned to a thickness of 4 µm and then subjected to H&E staining. Digital slide scanning was performed using the Panoramic P250 Digital Slide Scanner (3D Histech), as illustrated in Figure S2A.

### CODEX Workflow

The schematic overview of the CODEX workflow is depicted in Figure 1A. To validate the efficacy of commercial DNA-barcode-conjugated antibodies for PhenoCycler by Akoya Biosciences, we stained antibodies on tonsil tissue and aorta tissue microarray using the sample kit for PhenoCycler-Fusion (Akoya Biosciences, 7000017) following the manufacturer’s instructions. We tested 2 dilution factors (1:200, 1:500) for antibody titration. Staining patterns were confirmed using the online database, The Human Protein Atlas (www.proteinatlas.org).

**Figure 1.**
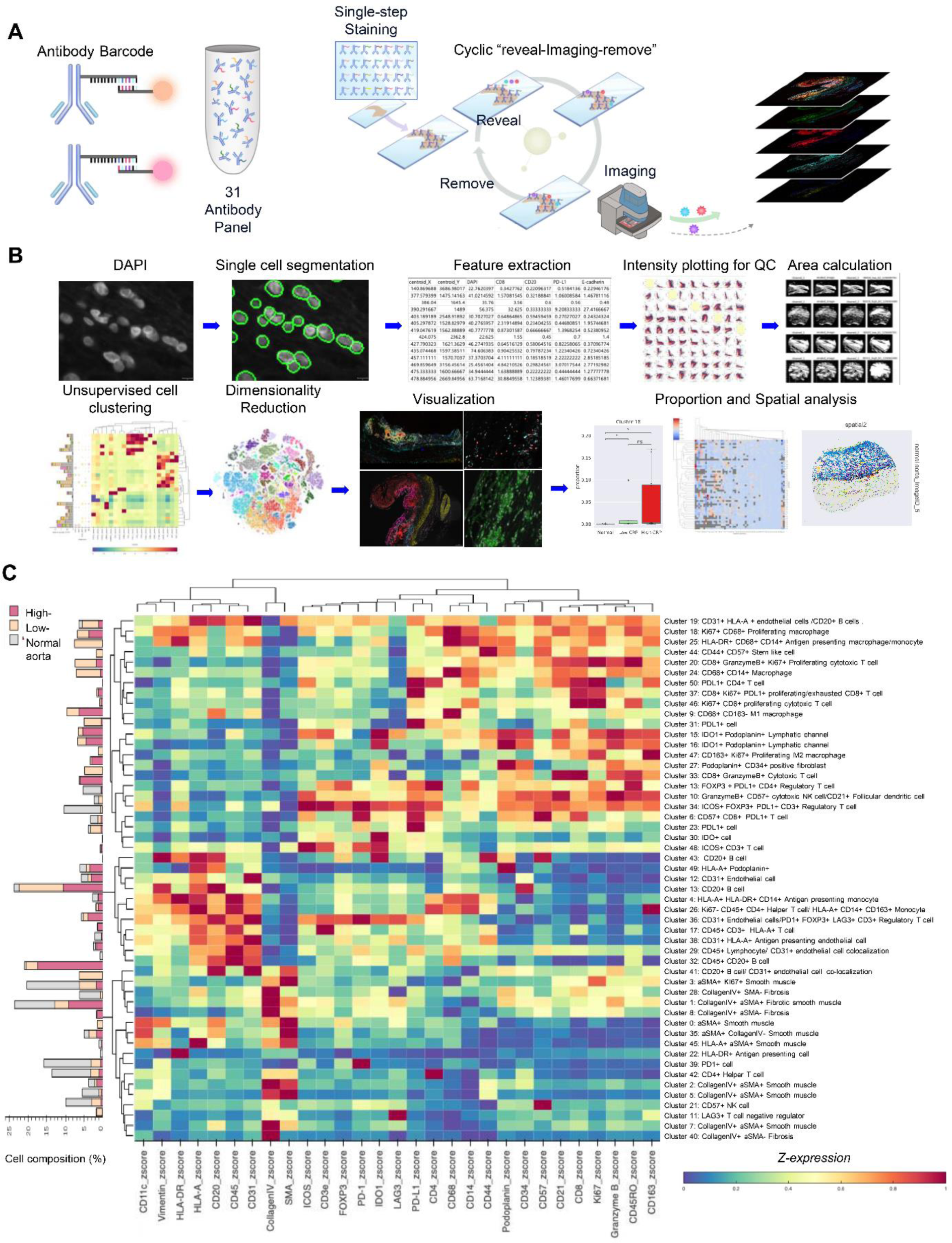
Defining single-cell phenotypes. (A) Schematic overview of the CODEX imaging workflow. We constructed 31 antibody panels tagged by oligonucleotides. All antibodies were stained in a single step, and imaging was performed on every three antibodies in a cyclic reveal-imaging-remove manner. All images were preprocessed and layered in one qptiff multidimensional image. (B) Computational analysis pipeline for multiplexed imaging data. After CODEX imaging, we segmented DAPI nuclei using Mesmer’s unsupervised segmentation method. Then, we extracted the mean intensity from each segmented cell, making 415,365 cells × 31 Antibodies matrix. Scatter plots for each antibody pair were visualized for quality control purposes. The area was calculated by binarized image for each ROI. Unsupervised clustering was performed for 415,365 cells using z score normalized mean intensity of 29 antibodies. We used the GPU version of Phenograph, Grapheno, for the unsupervised clustering. Fifty-one distinct phenotypes were identified. The right column contains narrative annotations for each clustering result. The heatmap was visualized by z expression score. Antibodies were also clustered, and same lineage antibodies were also clustered, which shows biological relevance. The left bar graph shows each cluster’s cell composition (%) of three groups: High-CRP, Low-CRP, and normal control. The composition ratio varies according to the phenotype cluster.

### Single-Cell Segmentation and Cell Mask Generation

The overview of the CODEX multiplexed imaging computational analysis workflow is illustrated in Figure 1B. Subsequently, we preprocessed the files and performed cropping for each TMA image. To effectively visualize multidimensional bioimaging data, we utilized the open-source visualization library Napari (https://github.com/napari/napari). To process large-sized, high-dimensional images, we utilized parallel computing techniques with the DASK Python library (https://www.dask.org).

We utilized the deep learning-based whole-cell segmentation algorithm MESMER^14^ to address the main challenges presented by the specimens, such as spindle cells, degenerative stroma, and atheromatous plaques (Figure S3). We used a DAPI signal for cell segmentation. We used np.stack to create a two-channel image and np.expand_dims to add an extra dimension to prepare the image for the segmentation algorithm. Upon visual inspection, we confirmed that the generated cell segmentation masks closely matched the tissue morphology. Given that a lymphocyte with a diameter of 7 µm corresponds to approximately 14 pixels, cells with an area larger than 400 pixels^2^ or smaller than 20 pixels^2^ were excluded from the analysis for biological relevance.

### Feature Extraction

We utilized the Scikit-image Python library (https://scikit-image.org) and its regionprops_table tool for image processing to extract the mean intensities from our CODEX high-dimensional image dataset. A total of 415,365 cells were processed to extract mean intensities for 31 antibodies, resulting in the creation of a cell matrix table with dimensions 415,365 by 31. We did not use the epithelial markers Pan-Cytokeratin and E-cadherin. A total of 29 antibodies were used for comprehensive phenotyping through unsupervised clustering.

Before conducting unsupervised clustering, we applied normalization procedures^15,16^. Cells were clipped at the 95^th^ percentile using the Pandas clip method (https://pandas.pydata.org/). Subsequently, we utilized the NumPy library’s arcsinh function to deskew the data. Following deskewing, Z normalization was performed using the formula below:

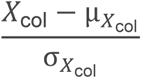

We used biaxial scatter plots to verify antibody intensities and observed that mutually exclusive markers display an L-shaped distribution (Figure S4). These normalization and validation steps were crucial in ensuring the accuracy and reliability of our data analysis, enabling robust downstream analyses and interpretation.

### Unsupervised Clustering and Phenotyping

We performed unsupervised clustering to discover novel cell clusters with previously unrecognized phenotypes. For this purpose, we utilized Grapheno^17^, an implementation of PhenoGraph (https://github.com/dpeerlab/PhenoGraph) with GPU acceleration using libraries such as CyPy^18^ and RAPIDS^19^. We set N_neighborhood to 40 and employed the Louvain community detection algorithm.^20^ As a result, the 415,365 cells were categorized into 51 distinct clusters (Figure 1C). To refine and characterize these clusters, extensive phenotype curation was carried out by a pathologist based on the z-values of each cluster. Out of the 51 identified clusters, those displaying strong expression of αSMA (α-Smooth Muscle Actin) and Collagen IV in a hierarchical manner, including clusters 0, 1, 2, 3, 5, 7, 8, 11, 21, 22, 28, 35, 39, 40, 42, and 45, were classified as stromal cells. The remaining clusters, which exhibited strong expression of various immune-related antibodies, were categorized as immune cells.

### Spatial Analysis: Neighborhood Enrichment Analysis

We conducted spatial neighborhood analysis using the open-source Python library Squidpy.^21^ To conduct spatial pattern and neighborhood enrichment analyses, we calculated the average shortest distance between reference cells and target cells. For neighborhood enrichment analysis, we used the “sq.gr.spatial_neighbors” and “sq.gr.nhood_enrichment” functions (https://squidpy.readthedocs.io/). This library calculates an enrichment score based on the proximity in the connectivity graph of cell clusters. The number of observed events is compared against permutations, and a z-score is computed. To visualize multiple images in a single cluster heatmap, we displayed the average cell-cell interactions for each image in the cluster map using the squidpy.pl.nhood_enrichment functions. We used open-source libraries Scanpy^22^ and the “sc.pl.spatial function” to apply pseudocolors to the clusters and perform in situ mapping (https://scanpy.readthedocs.io/).

### Statistical Analysis

For composition and density comparisons, we applied the two-sided Mann-Whitney-Wilcoxon test with Bonferroni correction for multiple comparisons at a significance level of 5% using the open-source Python library statannotations (https://pypi.org/project/statannotations), and SciPy (scipy.org).

More detailed descriptions of the methods can be found in the Detailed Methods section.

## RESULTS

### Clinical Characteristics of AAA and Normal Aorta

Baseline characteristics of patients with AAA or a normal aorta are shown in Table S2. In the normal aorta group, three patients underwent heart transplantation, and they showed elevated serum CRP levels before the operation; this was attributed to end-stage cardiomyopathies accompanied by multi-organ failure that necessitated mechanical circulatory support. However, patients with high serum CRP showed immunonegativity for CRP staining in their normal aorta.

### Multiplexed Staining of Aortic Tissues

By visualizing a total of 31 antibody-stained images using Napari, we observed significant differences in staining patterns among the normal aorta, Low-CRP, and High-CRP groups. In the normal aorta, immune cells and vascular structures were primarily distributed in the adventitia of the aortic wall. In contrast, in the case of AAA, immune cells were densely concentrated within the atherosclerotic plaque of the aneurysmal wall, and the expression patterns of various protein markers differed between the High-CRP group and the Low-CRP group within abdominal aortic aneurysms (Figure 2). Immune cells, including CD68-positive macrophages, were focally aggregated in the adventitia of a normal aorta. In contrast, the Low-CRP group exhibited intense immune marker staining in the atherosclerotic plaque in the vascular media, along with fibrosis. Furthermore, in the High-CRP group, diffuse and intense inflammatory cell infiltration was identified in atherosclerotic plaques mainly deposited on the intima of the vascular wall (Figure 2C). By visualizing several antibodies with multiple layers, we confirmed that the composition of immune cells differed between the Low-CRP and High-CRP groups (Figure 2D).

**Figure 2.**
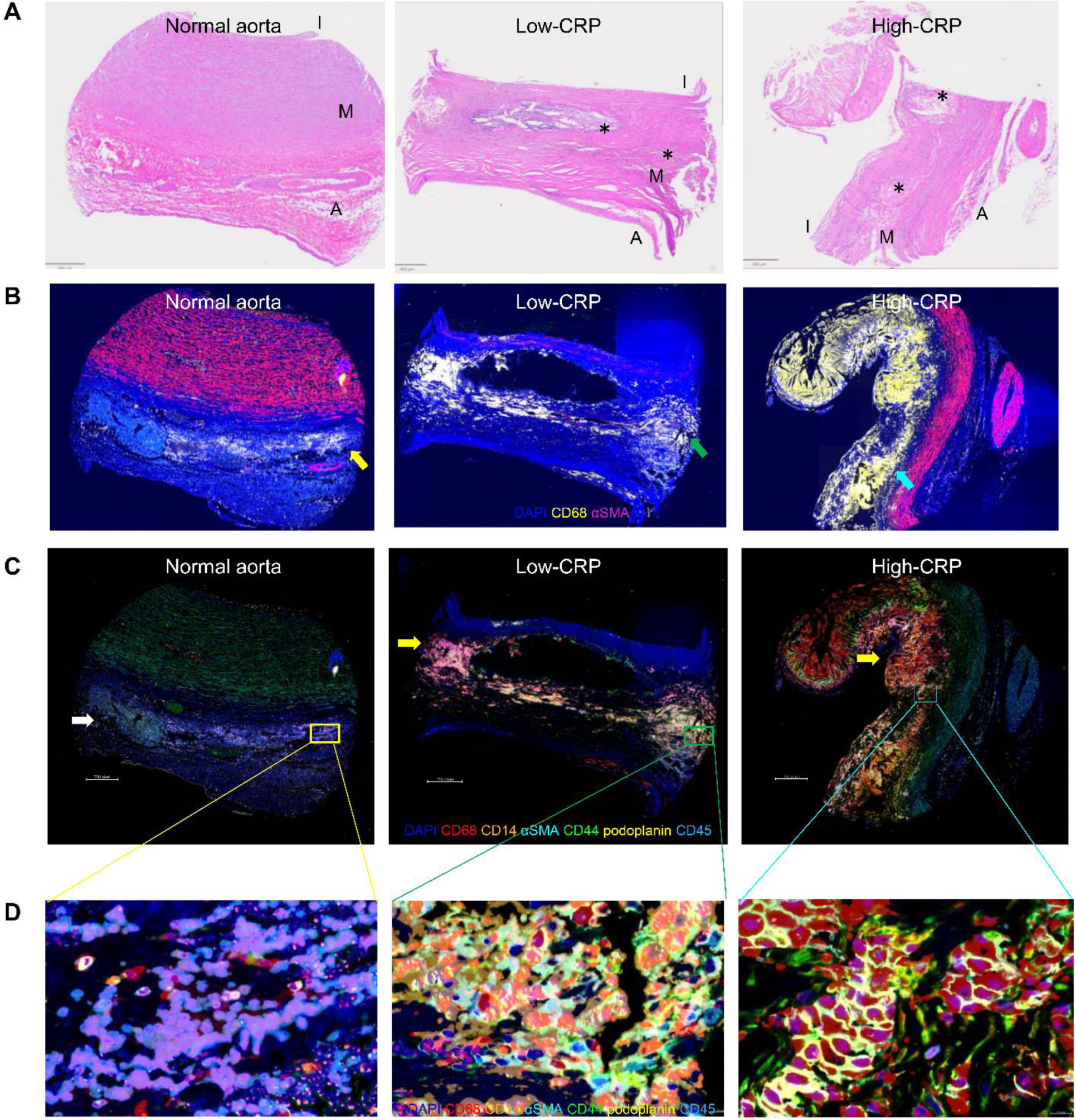
Multiplexed staining images for normal aorta, Low-CRP, and High-CRP. (A) Corresponding H&E staining images to multiplexed images of B-D of normal aorta, Low-CRP, and High-CRP. I, intima; M, media; A, adventitia; * Atherosclerotic plaque (B) Visualization of DAPI (blue), smooth muscle actin (magenta), and CD68 (yellow) staining. CD68-positive macrophages were focally aggregates in the adventitia of normal aorta (white arrowhead) and atherosclerotic plaque in media along with fibrosis of Low-CRP (green arrowhead). Diffuse and intense inflammatory cell infiltration was identified in atherosclerotic plaque of High-CRP (cyan arrowhead). (C) Overlayed image of multiplexed six antibodies and DAPI. In AAA, intense immune cell infiltration was identified in atherosclerotic plaque in intramural atherosclerotic plaque and atherosclerosis, replacing intima of the aorta in High-CRP. (D) Magnified view of multiplexed imaging. Different antibody staining patterns were highlighted in low-CRP and high-CRP.

### Cell Phenotyping Using an Unsupervised Clustering Algorithm

We successfully segmented 415,365 cells using a deep learning-based approach for cell segmentation. We calculated the mean intensity values of 31 different antibody channels for these cells, resulting in a 31 x 415,365 mean intensity table for the analysis. Subsequently, we performed extensive normalization on these cells and mean intensity values using arcsine transformation and z-score normalization. Because there were no epithelial cells in our sample, we excluded E-cadherin and pan-cytokeratin, which are epithelial markers used for phenotyping. Through unsupervised clustering, we identified fifty-one distinct cell clusters (Figure 1C). Markers with relevant biological functions were clustered together, indicating the robustness of identifying biologically meaningful cell types using unsupervised clustering in Grapheno. For example, the clusters HLA-DR/HLA-A as antigen presentation markers, and Collagen IV/αSMA were clustered as stromal markers, while FOXP3/PD-1, IDO-1, LAG3, and PD-L1 were clustered as immune regulators, thereby supporting the notion that the unsupervised clustering was appropriately performed.

We performed t-SNE dimensionality reduction to embed all cells, and the t-SNE plots showed that each phenotype formed well-defined clusters (Figure 3A). Additionally, the t-SNE plots revealed distinct clusters not only between the normal aorta and AAA but also among different CRP levels within AAA (High-CRP vs. Low-CRP) (Figure 3B). We visualized the mean intensity z-scores of each marker on the t-SNE plot. Markers of different biological functions occupied different clusters that were well separated, indicating that the clustering results are biologically relevant overall (Figure S5).

**Figure 3.**
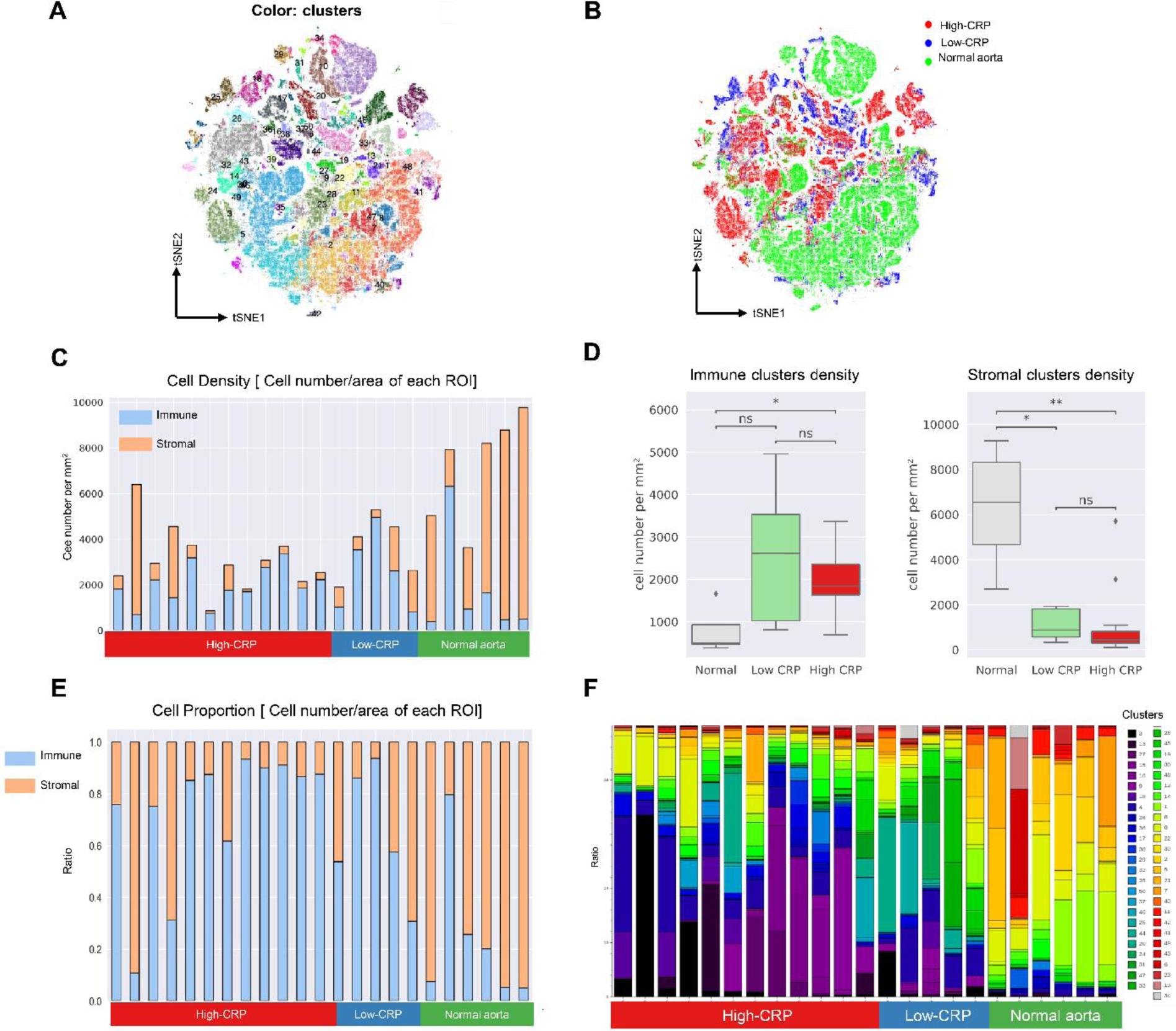
Comparison of phenotype composition between High-CRP, Low-CRP, and normal aorta. (A) In t-SNE plots, phenotype clusters were overlayed with different colors. Each phenotype cluster was aggregated in t-SNE, showing the reliable clustering result by unsupervised clustering, Grapheno. (B) In t-SNE plots, study groups with CRP levels were overlayed with different colors. According to the presence of AAA and CRP levels, the plots were well divided into t-SNE plots. (C) According to the study group, Immune and stromal cell density [cell number/mm^2^, Immune or stromal cell number/area of each ROI]. As shown in (D), immune cell density was higher in AAA with Low-CRP and High-CRP groups, and stromal cell density was lower in AAA with Low-CRP and High-CRP groups. *; p-value < 0.05, **; p-value <0.005. ROI; Region of interest (E) Lineage proportion comparison. Upon analyzing the cell proportion by dividing the total number of cells in each ROI, it was found that the AAA group exhibited a higher percentage of immune cells relative to the overall cell population. (F) Cell phenotype cluster composition according to the CRP level. The composition of cell phenotypes varied among the High-CRP, Low-CRP, and normal aorta groups.

### Decrease of Stromal cells and Enrichment of Immune cells in AAA

When we performed hierarchical clustering based on the expression of 29 markers, specifically focusing on collagen IV and αSMA, which are two main vascular stromal markers, we observed a distinct separation into two major clusters that were predominantly composed of stromal cells (clusters 0, 1, 2, 3, 5, 7, 8, 11, 21, 22, 28, 35, 39, 40, 42, 45) and the rest of the clusters for immune cells.

Upon comparing the absolute cell numbers per TMA core, we observed a significant decrease in the total cell count and cell density in the aortic aneurysm compared to the normal aorta (Figure 3C). This decrease was particularly noticeable in the stromal cell population among the total cell population, and the immune cell proportion was higher in AAA than in the normal aorta (Figure 3D)

### Twenty-four Clusters enriched in AAA

A total of 20 clusters (clusters 9, 15, 16, 18, 20, 24, 25, 27, 31, 33, 37, 44, 46, 47, 50), highlighted in Figure 1C, were completely absent in the normal aorta. Twenty-four specific cell clusters were more frequently observed in the AAA-high or Low-CRP groups (clusters 2, 4, 5, 7, 9, 11, 15, 16, 17, 18, 20, 24, 25, 26, 28, 33, 36, 38, 39, 40, 41, 46, 47, 49). According to the CRP level, the composition of the specific phenotype cluster was different (Figure 1C, left bar plot, and Figure 3F). These observations suggest the presence of unique immune and stromal microenvironments in AAA, which are further influenced by the CRP level. In situ visualization and pseudo-mapping for each cluster showed that the immune cell environment markedly differed between the normal aorta, High-CRP, and Low-CRP groups (Figure 4).

**Figure 4.**
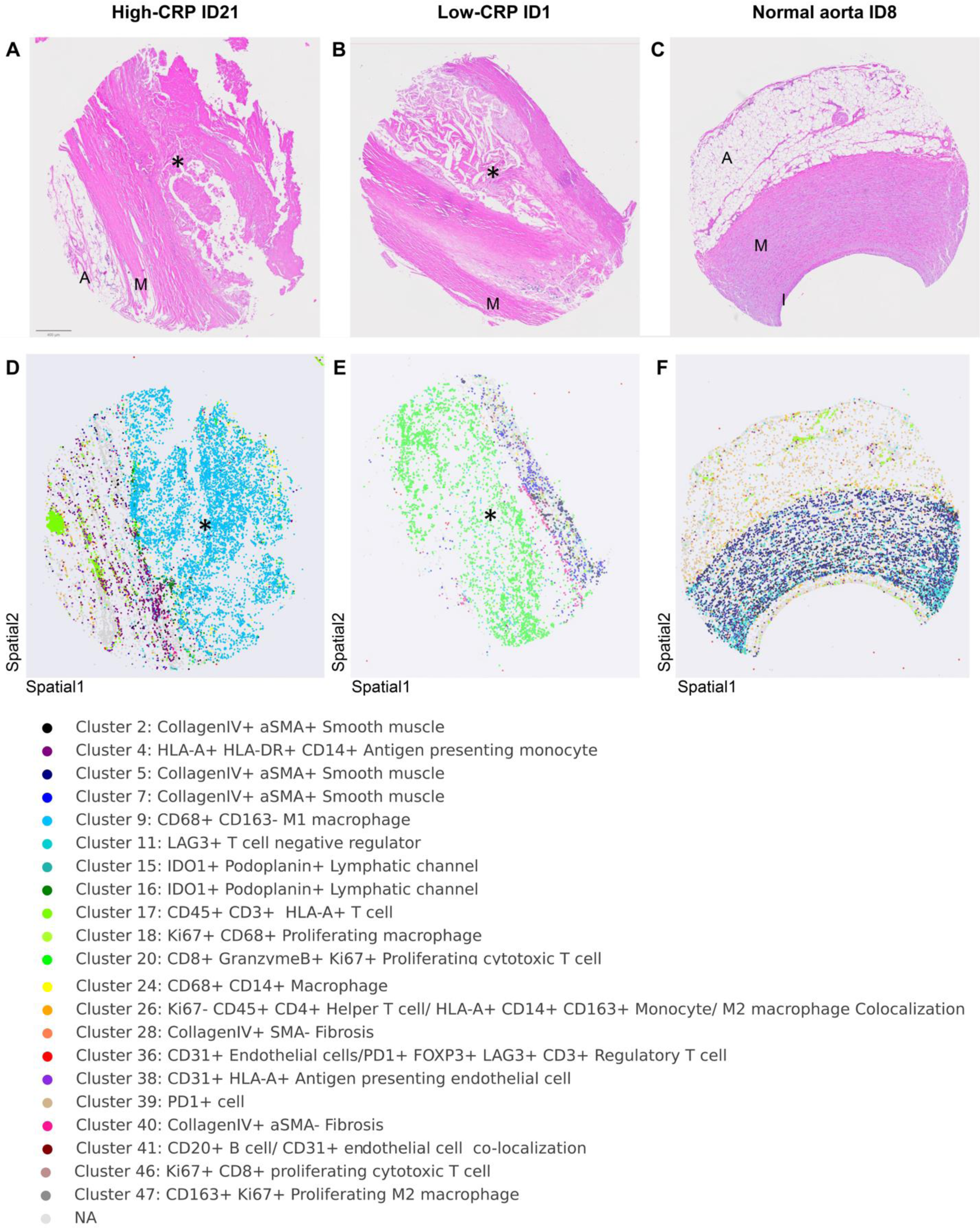
In situ visualization of pseudo-mapping for each cluster using Squidpy. Among the 51 phenotype clusters, the representative clusters that showed the different cell phenotype composition between AAA vs normal aorta were visualized. The cell phenotypes of the atheroma varied significantly depending on the CRP level. (A-C) Corresponding H&E staining images to D-F cell phenotype map of High-CRP, Low-CRP, and normal aorta. (D-F) A cell phenotype map showing cell phenotype clusters by color. (I, intima; M, media; A, adventitia; * Atherosclerotic plaque)

### Cell Composition Comparison

To compare the composition of specific clusters across individual cores, we normalized the cell counts of these clusters by dividing them by the total number of immune or stromal cells in each core. Specifically, for clusters identified as immune cells, we divided the cell count by the total number of immune cells in each core. For clusters corresponding to stromal cells, we divided the cell count by the total number of stromal cells in each core. In the case of AAA, we observed a consistent trend, particularly within the high CRP group, where the proportion and density of stromal-immune cell clusters displayed similar patterns of variation (Figures S6, S7).

AAA groups showed an enrichment of CD3+ T cells (cluster 17) along with a decrease in LAG3+ cells (cluster 11), which function as immune checkpoints and negatively regulate T cell proliferation, activation, and T cell homeostasis. On the contrary, LAG3+ cells were abundant in the walls of normal aortic vessels. This suggests that T cell activation may be more enhanced in aneurysms associated with a reduction in negative regulators of T cells such as LAG3^23^. Furthermore, the proliferation of CD3-positive T cells (cluster 17) was notably higher in the high CRP subgroup, and this phenotype was found in the atheromatous core (Figure 5, A and B).

**Figure 5.**
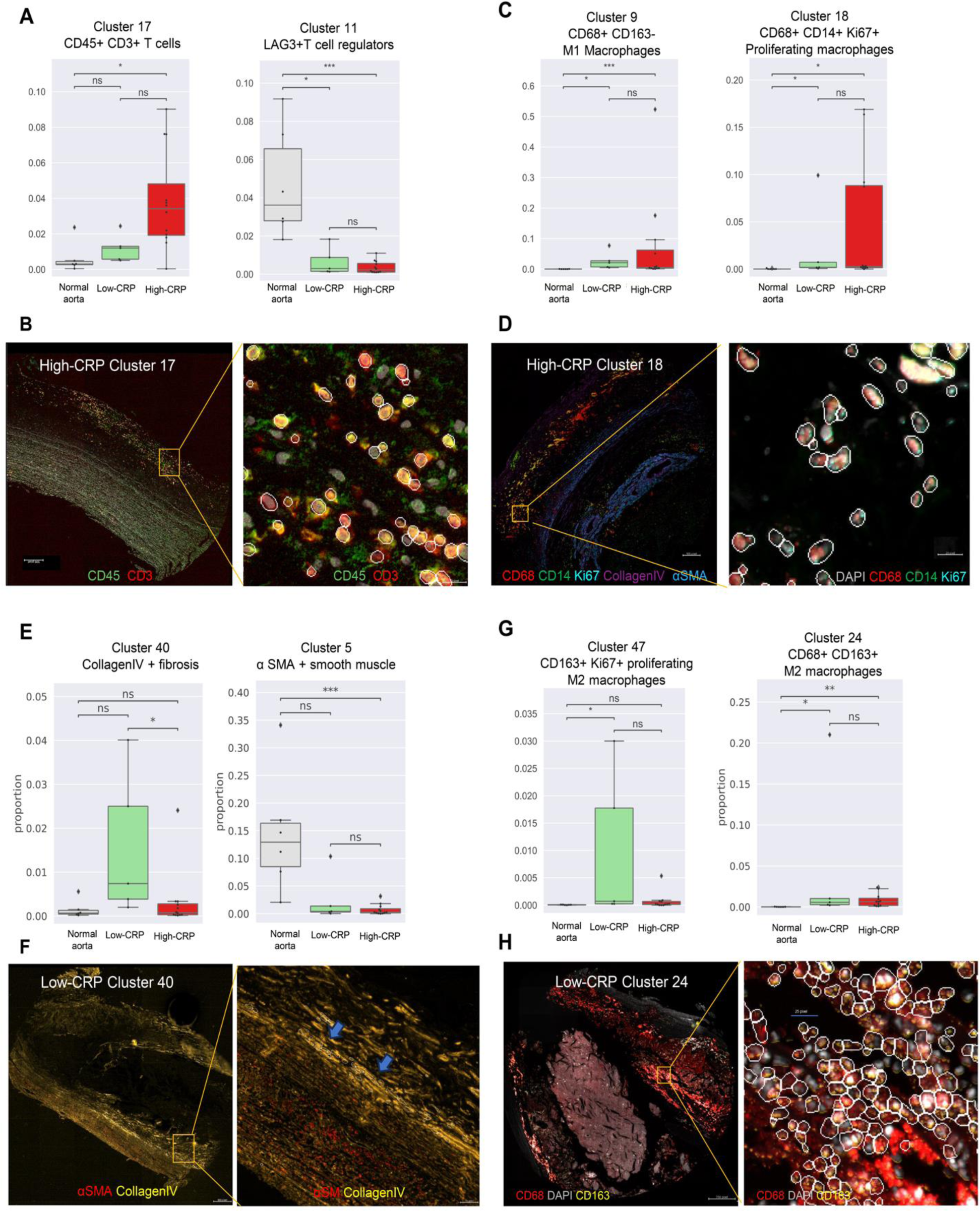
Cluster proportion comparison between High-CRP, Low-CRP, and normal aorta, and visualization of multiplexed antibodies. (A) Cluster 17, consisting of CD45+CD3+ T cells, increased in AAA, particularly in those with high CRP. This trend contrasted with the proportion of cluster 11, which is a LAG3+ T cell negative regulator. (B) Within the atherosclerotic plaque of High-CRP (orange box), cells of cluster 17 were visualized using antibodies against CD45 and CD3, demonstrating co-localization of the two antibodies. (C) The cell proportions of cluster 9, which consists of CD68+ CD163-M1 macrophages, and cluster 18, identified as CD68+ CD14+ Ki67+ proliferating macrophage/monocytes, significantly increased in High-CRP. (D) Representative image of Cluster 18 observed within the atheromatous plaque of High-CRP (indicated by an orange box). (E) Cluster 40, characterized by Collagen IV+ fibrosis, was markedly expressed in the Low-CRP and was identified in the remaining thinned aortic wall. Alpha-smooth muscle actin-positive (αSMA+) smooth muscle cells, represented by cluster 5, significantly decreased in cases of aortic aneurysm. (F) Representative image of Cluster 40 observed in the fibrotic vascular wall in AAA with Low-CRP (orange box). αSMA-Collagen IV+ fibrosis was visualized with two antibodies. (G) CD163+ Ki67+ proliferating M2 macrophages increased significantly in the Low-CRP group, exhibiting a pattern similar to that of cluster 40, which is associated with Collagen IV+ fibrosis. CD68+ CD163+ M2 macrophages were increased in AAA. (H) The expression of CD68+ CD163+ M2 macrophages was observed within the atheromatous plaque of the Low-CRP group (orange box). (Indicated specific phenotype cluster was highlighted by a white line on the cell segmentation mask outline.)

Compared to a normal aorta, the AAA groups exhibited an increase in CD68+CD163-M1 macrophages (cluster 9) and CD68+ CD14+ Ki67+ (cluster 18) proliferating macrophages/monocytes, which was more pronounced in the high CRP group (Figure 5C). Also, these macrophage phenotypes were identified within the core of the atherosclerotic plaque (Figure 5D).

When comparing the high and low CRP subgroups, we observed a significant increase in fibrosis (collagen IV+ αSMA-; cluster 40) within the vascular wall (Figure 5, E and G) in the Low-CRP group. This trend was also found in the proliferating M2 macrophage clusters characterized by CD163+ Ki67+ expression (cluster 47, Figure 5G). Simultaneously, AAA with both low and high CRP showed an increase of CD68+ CD163+ M2 macrophage clusters (cluster 24, Figure 5, G and H).

In the low CRP group, there was a relative increase in the stromal portion within the immune/stromal ratio, suggesting that M2 macrophages are likely to be associated with fibrosis in AAA. Furthermore, we observed a decrease in αSMA-positive normal vascular smooth muscle cells in AAA. Particularly within the high CRP subgroup, clusters 2 and 7, characterized by collagen IV+αSMA+ cells distributed within the vessel media, exhibited a more significant reduction. This suggests that there is a depletion of normal smooth muscle cells in AAA, which is more pronounced in the high CRP group (Figure S6, S7).

### Neighborhood Enrichment Analysis

We applied neighborhood enrichment analysis^21,24^ to identify non-randomly distributed cells of a specific phenotype in relation to another phenotype within a tissue sample using the spatial coordinates of each cell. This method compares the observed events against a random configuration to assess the non-random distribution of phenotype clusters. This analysis provides information on the interactions of different cell clusters within a tissue^25,26^. Then, we calculated the spatial proximity by determining the shortest average distance between each cell in two phenotype clusters identified from the neighborhood enrichment analysis heatmap, which displayed statistical significance. The enrichment patterns varied based on the CRP level among AAA (Figure 6).

**Figure 6.**
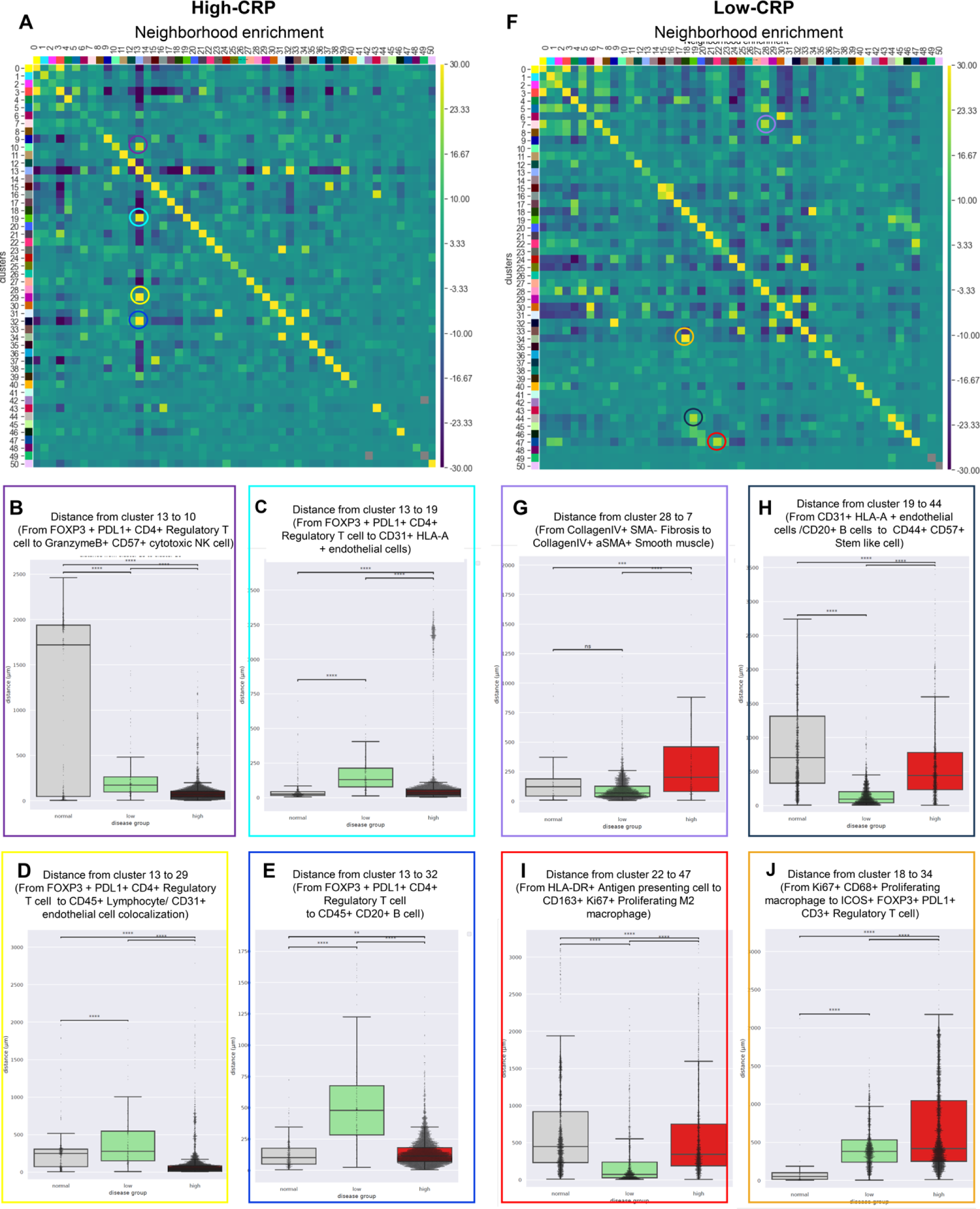
Neighborhood enrichment analysis and spatial distance comparison. (A) Neighborhood enrichment analysis of High-CRP. In High-CRP, several phenotype clusters (clusters 10, 19, 29, 32) were significantly enriched in proximity to cluster 13 (FOXP3+ PDL1+ CD4+ Regulatory T cells). (B-E) Shortest average distance comparison from one cell phenotype cluster to another in High-CRP. One dot indicates the cell. Distance from cluster 13 (FOXP3+ PDL1+ CD4+ Regulatory T cells) to cluster 10 (GranzymeB+ CD57+ cytotoxic NK cell/CD21+ Follicular dendritic cell), cluster 19 (CD31+ HLA-A + endothelial cells /CD20+ B cells), cluster 29 (CD45+ Lymphocyte/ CD31+ endothelial cell colocalization), cluster 32 (CD45+ CD20+ B cell) were significantly shorter in High-CRP. (F) Neighborhood enrichment analysis of Low-CRP. In Low-CRP, different clusters were enriched compared to the High-CRP. (G-J) Shortest average distance comparison from one cell phenotype cluster to another in Low-CRP. Distance from cluster 28 to 7 (From CollagenIV+ αSMA-Fibrosis to CollagenIV+ αSMA+ Smooth muscle), from cluster 19 to 44 (From CD31+ HLA-A + endothelial cells /CD20+ B cells to CD44+ CD57+ Stem-like cell), from clusters 22 to 47 (From HLA-DR+ Antigen presenting cell to CD163+ Ki67+ Proliferating M2 macrophage), and from cluster 18 to 34 (From Ki67+ CD68+ Proliferating macrophage to ICOS+ FOXP3+ PDL1+ CD3+ Regulatory T cell) were significantly shorter in Low-CRP. (Statistics for B-E, and G-J: Mann–Whitney–Wilcoxon test two-sided with Bonferroni correction: ns (not significant): p > 0.05; *: p <=0.05; **: p <=0.01, ***: p <=0.001, ****: p <=0.0001. Color around the box plot of the shortest average distance comparison corresponds to the color box in the neighborhood enrichment analysis heatmap in A and F.).

In the High-CRP group, several phenotype clusters (clusters 10, 19, 29, 32) were significantly enriched in proximity to cluster 13 (FOXP3+ PDL1+ CD4+ regulatory T cells). By comparing the shortest average distances, we found that the distances from cluster 13 (FOXP3+ PDL1+ CD4+ regulatory T cells) to cluster 10 (granzyme B+ CD57+ cytotoxic NK cell/CD21+ follicular dendritic cell), cluster 19 (CD31+ HLA-A + endothelial cells /CD20+ B cells), cluster 29 (CD45+ lymphocyte/ CD31+ endothelial cell colocalization), and cluster 32 (CD45+ CD20+ B cells) were significantly shorter in the High-CRP group.

The Low-CRP group showed a different neighborhood enrichment pattern compared to the High-CRP group (Figure 6F). Shortest average distances from cluster 28 to 7 (from collagen IV+ αSMA-fibrosis to collagen IV+ αSMA+ smooth muscle), from cluster 19 to 44 (from CD31+ HLA-A + endothelial cells/CD20+ B cells to CD44+ CD57+ stem-like cell), from clusters 22 to 47 (from HLA-DR+ antigen-presenting cells to CD163+ Ki67+ proliferating M2 macrophages), and from cluster 18 to 34 (from Ki67+ CD68+ proliferating macrophages to ICOS+ FOXP3+ PDL1+ CD3+ regulatory T cells) were significantly shorter in the Low-CRP group.

## DISCUSSION

This study is the first to utilize multiplexed imaging to investigate the immune-stromal response in the pathophysiology of AAAs. We analyzed the immune and stromal microenvironment in relation to CRP immunodeposition in atheromatous plaques, incorporating spatial information. In addition to the well-documented processes of aortic wall thinning, intima erosion, and atheroma formation in AAA pathophysiology, we observed a significant increase in the presence of immune cells and alteration of immune cell phenotype, providing evidence of a robust immune response in AAA, especially with CRP deposition. Simultaneously, there were also notable fibrotic changes in the stromal cell component in the Low-CRP group, suggesting that there might be differences in the pathophysiologic fibrotic changes according to the degree of CRP deposition in AAA. We visualized multiple antibodies simultaneously on atheroma, enabling a more direct and intuitive confirmation of the immune cell environment within atheroma of AAA. This approach provided strong evidence suggesting that atherosclerosis-related AAA is indeed associated with inflammatory processes, further highlighting the significance of inflammation in the context of AAA. Importantly, we previously found that as serum CRP levels increased^10^, the degree of CRP deposition within atheroma also increased, supporting the active and critical role of CRP in the pathogenesis of aortic aneurysms.

Although we excluded cases with inflammatory processes such as chronic aortitis, we found that atheroma contains numerous inflammatory cells representing various immune subtypes. There was a shift towards proinflammatory M1 macrophages and an increase in monocyte subtypes involved in antigen presentation in AAA, particularly in cases with high levels of CRP. Conversely, in the Low-CRP group, immune cells shifted towards M2 macrophages, leading to an increase in fibrosis. These results suggest that AAA can be classified into two subtypes based on serum CRP levels and CRP deposition. This classification leads to distinct immune cell compositions and immune response patterns within the aneurysm.

Our study aligns with the results of in vitro research conducted by Devaraj *et al*.,^27^ which showed that when monocytes were incubated with CRP in an atherosclerotic environment, macrophages tended to polarize towards the M1 phenotype. In contrast, the transition to the M2 phenotype was inhibited. The present study has confirmed similar results in surgically resected specimens from patients with AAA, providing in situ evidence that supports previous in vitro research findings. Monocytes play a critical role in the immune system as control switches that regulate the balance between pro-inflammatory and anti-inflammatory activities. Furthermore, accumulating studies have proven that the M1/M2 polarization state of circulating monocyte-derived macrophages plays a crucial role in regulating the development of AAA^28^. In addition, targeting M1/M2 macrophage polarization has been proposed as a potential strategy to control inflammation and the progression of AAA. Pope et al. showed that D-series resolvins could inhibit the formation of murine AAA by promoting M2 macrophage polarization^29^. Dale et al. reported that elastin-derived peptides promote AAA formation by modulating M1/M2 macrophage polarization^30^. In addition, our previous study showed that AAA with intense CRP deposition was related to the poor prognosis of aortic aneurysm^10^. n line with these studies, the present study suggests that targeting M1/M2 macrophage polarization associated with CRP deposition could be a novel molecular approach to impede AAA progression.

The observation that atheroma in the high CRP group exhibited a complex multicellular spatial organization with a diverse composition of immune and stromal cells aligns with recent findings in the field of aortic aneurysms. Notably, a study by Cao *et al.*^31^ investigated the phenotypic changes in various vascular smooth muscle cells using single-cell RNA sequencing and demonstrated that vascular walls undergoing phenotypic switches to fibroblast-like vascular smooth muscle cells (VSMCs) exhibited increased collagen secretion and differentiated into T-cell-like VSMCs and macrophage-like VSMCs. Via unsupervised clustering for phenotyping, we discovered an unpredicted novel phenotype cluster 42, which showed weak Collagen IV expression, weak αSMA expression, and strong CD4 expression within the stromal cluster. Further study is needed if this novel phenotype corresponds to the Cao’s VSMCs.

The JUPITER study,^32^ which is an intervention trial evaluating the use of rosuvastatin, has shown that administering rosuvastatin preventively to individuals with serum hs-CRP levels of 2 mg/L or higher, even in the absence of other risk factors, resulted in a significant reduction in cardiovascular events. A recent systematic review and meta-analysis have highlighted the role of statins in lowering serum CRP levels in patients with cardiovascular disease.^33^ Furthermore, several studies have shown that statin medication was effective not only in halting the progression of AAA, but also in decreasing the risk of rupture by suppressing inflammatory mediators^34^, and a meta-analysis showed that statin therapy may be associated with a reduction in AAA progression and rupture^35^. These findings on CRP levels, the effects of CRP-lowering statins, and the therapeutic impact of statins on AAA progression collectively support the notion that targeting CRP in AAA can significantly influence the development of aneurysms as a novel therapeutic target.^36^

LAG3 (Lymphocyte activation gene-3), also known as CD223, is a negative regulator of T-cell activation and plays a crucial role in modulating the immune response. LAG3 is required for optimal T cell regulation and homeostasis^23,37^ and it is receiving significant attention as an immune checkpoint modulator and a new therapeutic target for anti-cancer immunotherapy^38^. LAG3 is not only expressed on multiple cell types, mainly in CD4+ and CD8+ T cells, but LAG3 RNA expression was also identified in smooth muscle and heart, as shown in the Human Protein Atlas (https://www.proteinatlas.org/ENSG00000089692-LAG3/tissue), and aorta regulatory T cells showed a high expression level of Treg signature genes, including Lag3^39^. Our study showed that LAG3 was expressed in the normal aortic vascular wall but was absent in atherosclerotic plaque, along with an accumulation of CD3+ lymphocytes in the atherosclerotic plaque. In line with our study, Mulholland et al. reported that CD4+ T cell accumulation in aortic plaque was observed in LAG3 knock-out mice and mice treated with LAG3 blockade treatment^40^. Further study of the effect of LAG3 on atherosclerosis with aortic aneurysm is needed.

Our study demonstrates the feasibility of multiplexed imaging techniques such as CODEX, even when using over a decade-old FFPE blocks of collagen-rich vascular tissue, for detecting significant signals. This result highlights the potential for conducting retrospective research on vascular diseases, including rare cases, using archived tissue samples with FFPE for newly developed multiplexed imaging technology. This approach will unlock numerous research opportunities for analyzing historic cardiovascular specimens related to various immune responses that have been stored but not previously studied.

It is also noteworthy that we utilized unsupervised clustering techniques, which enabled us to discover cluster phenotypes that were previously unknown. Supervised clustering based on known phenotypes is often conducted in many high-dimensional multiplexed imaging studies. However, this approach is limited to confirming only the phenotypes that are already known. Using unsupervised algorithms, such as PhenoGraph^41^ accelerated by GPU via RAPIDS, we conducted clustering without relying on prior biological knowledge. This approach precisely identified known cell types with high cluster coherence, minimizing bias introduced by prior biological knowledge.^42^

The unsupervised clustering technique allowed us to identify Ki67-positive proliferating macrophages, a cell population that is not commonly addressed in routine immune cell phenotyping. Ki67 is a universally applicable marker in cell proliferation, expressed throughout all active phases of the cell cycle. While macrophages are typically considered terminally differentiated cells without mitotic activity, recent studies have reported the presence of proliferating macrophages expressing Ki67, indicating self-renewal capabilities. Lhotak *et al.* have described the presence of Ki67+ CD68+ proliferating macrophages in coronary artery atherosclerosis,^43,44^ suggesting their role in initiating and sustaining inflammation.^45^ Further investigations are warranted to explore the metabolic and epigenetic reprogramming associated with macrophage polarization in atherosclerotic AAA. Furthermore, targeting macrophage polarization presents a novel therapeutic opportunity for reducing residual inflammation and addressing cardiovascular diseases.

The tissue samples containing atheroma exhibited high levels of autofluorescence, which posed a challenge for CODEX, a fluorescence-based imaging modality. This resulted in nonspecific signals that hindered the direct incorporation of CRP into supervised clustering. As the size of our cohort was rather small, more samples might be needed to further validate the discoveries with stronger statistical power.

In conclusion, we utilized CODEX to precisely map the multicellular dynamics of aortic aneurysms based on the level of CRP deposition. Our findings suggest a potential association between the immune landscape related to CRP and the progression of aortic aneurysms. Targeting these specific immune cell components offers a promising and innovative therapeutic strategy to mitigate the progression of aortic aneurysms. Further research and clinical investigations are needed to validate these findings and explore the full therapeutic potential of targeting these immune cell components in managing aortic aneurysms.

## Non-standard Abbreviations and Acronyms

αSMA: α-Smooth Muscle Actin
AAA: Abdominal aortic aneurysm
CRP: C-reactive protein
CODEX: Co-detection by indexing
FFPE: Formalin-fixed paraffin-embedded
H&E: Hematoxylin and eosin
IHC: immunohistochemistry
IRB: Institutional Review Board
LAG3: Lymphocyte-activation gene 3
mCRP: Monomeric CRP
pCRP: pentameric form of serum CRP
ROI: Region of Interest
TMA: Tissue microarray
t-SNE: t-distributed stochastic neighbor embedding
VSMC: Vascular smooth muscle cell

## Acknowledgments

We thank all the patients who contributed to the study. We thank Kyung Min Park, Gil Je Lee, and Jong Man You for the setting up the CODEX multiplexed imaging system in Korea.

## Supplemental Material

### Detailed Methods

**Table S1**. Dilution Ratio and Volume Requirement of Antibodies for Staining Solution

**Table S2**. Patient characteristics

**Figure S1**. Representative anti-CRP antibody immunohistochemical staining of the High-CRP (left) and Low-CRP (right). This anti-CRP antibody detects both pentameric CRP and monomeric CRP. The High-CRP group showed diffuse and strong immunopositivity for the anti-CRP antibody, while the Low-CRP group showed negativity for the anti-CRP antibody.

**Figure S2. Scan image of Tissue microarray (TMA)**

(A) Digitalized image of hematoxylin and eosin staining

(B) Scan view of CODEX multiplexed imaging

**Figure S3**. Cell segmentation mask visualization on normal aortic wall.

**Figure S4**. Signal intensity scatter plot.

We performed scatter plots for antibodies known to exhibit mutually exclusive expression patterns. This set of antibodies formed an L-shaped plot. Additionally, epithelial markers were negative in vascular samples. These findings indicate the proper staining of the antibodies.

**Figure S5**. In the t-SNE plot, the normalized expression of each marker was overlaid.

**Figure S6**. Cell proportion comparison. Twenty-four specific cell clusters (clusters 2, 4, 5, 7, 9, 11, 15, 16, 17, 18, 20, 24, 25, 26, 28, 33, 36, 38, 39, 40, 41, 46, 47, 49) were significantly more frequently observed in the AAA-high or Low-CRP groups.

**Figure S7**. Comparison of cell density [cell number/mm^2^, Immune or stromal cell number/area of each ROI] in normal aorta, Low-CRP, and High-CRP groups. The trend was similar to the composition comparison graph in Figure S6.

**Figure S8**. Creation of binary images and calculation of area through αSMA image modification.

