## Supplemental Material for "Unraveling the Complexity of Abdominal Aortic Aneurysm: Multiplexed Imaging Insights into C-Reactive Protein-Related Variations"

### Detailed Methods

#### Ethics Statement

The study protocol was approved by the Institutional Review Boards (IRB) of Asan Medical Center (IRB number: S2020-0196) and SMG-SNU Boramae Medical Center (IRB number: 20-2022-111). The study protocol adhered to the relevant ethical guidelines and regulations. Because the bioethical law in South Korea mandates the need for written informed consent when using biological specimens obtained since 2013, we conducted multiplexed imaging CODEX staining using formalin-fixed paraffin-embedded (FFPE) tissue blocks obtained in 2012 with anonymized patient information. Both IRBs approved using clinical data and collecting and utilizing biological samples for research purposes, waiving the need for formal written informed consent.

#### Histopathological Assessment and Immunohistochemical Staining

The excised aortic tissue was sectioned into 5 mm intervals for the pathological examination of resected aortic aneurysms. Visual inspection was conducted to identify areas with atheromatous plaque deposition and thinning of the arterial wall. Tissue sections that met these criteria, including a minimum of three regions, were preserved in 10% buffered formalin for one day to prepare FFPE blocks.

Unstained tissue slides (thickness: 4 µm) were prepared from the FFPE blocks. Hematoxylin and eosin (H&E) staining was performed for histopathological evaluation. Immunohistochemical staining was conducted using anti-CRP antibodies (rabbit polyclonal, ab32412, Abcam, Cambridge, UK) capable of detecting both the monomeric and pentameric forms of CRP. Additionally, anti-mCRP antibodies (mouse monoclonal, C1688, Sigma-Aldrich, Saint Louis, MO, USA), which selectively detect monomeric CRP, were used to stain the tissue sections. Because the results of CRP immunostaining and mCRP immunostaining were well correlated[11], we used the CRP immunostaining results for patient grouping.

**FFPE Tissue Pre-Treatment and Antibody Staining**

TMA sample slides were initially incubated at 65°C for 1 hour to melt the paraffin. Then, tissue deparaffinization and hydration were performed in the following order for 5 minutes each: xylene, xylene, 100% ethanol, 100% ethanol, 90% ethanol, 80% ethanol, 70% ethanol, 50% ethanol, 30% ethanol, $ddH_2O$, and $ddH_2O$. Next, the sample slide was fixed with a 4% paraformaldehyde solution in PBS (T&I, BPP-9004) for 1 hour and washed with ddH2O for 5 minutes twice before proceeding to antigen retrieval.

The stock 10x AR9 Solution (Akoya Biosciences, #AR9001KT) was diluted to 1x with $ddH_2O$ for antigen retrieval. The sample slide was then placed in the vessel with 250 mL of 1x AR9 buffer, ensuring that the slide was fully submerged in the pressure cooker. The cooker was set to high pressure (11.6 PSI/110°C) for a cooking duration of 20 minutes. Because our samples contain atheroma that can be easily dethatched, we substituted heating with the conventional water bath during antigen retrieval to solve this issue. Once the run was complete, the sample slide was removed from the cooker and allowed to cool down at room temperature for 1 hour. After cooling to room temperature, the sample slide was washed with $ddH_2O$ and then incubated for 2 minutes. Next, the sample slide was incubated with the

Hydration Buffer (Akoya Biosciences, #7000017) for 2 minutes and repeated. The sample slide was then incubated with the Staining Buffer (Akoya Biosciences, #7000017) for 20 minutes to allow the sample to reach equilibrium.

**FFPE Tissue Staining**

The antibody cocktail stock solution was prepared with 31 antibodies for cell lineage markers, cell status, and immune checkpoint proteins. The antibody panel was as follows: CD8-BX026 (Akoya Biosciences, #4250012, Marlborough, MA, USA), E-cadherin-BX014 (Akoya Biosciences, #4250021), CD14-BX037 (Akoya Biosciences, #4450047), Ki67-BX047 (Akoya Biosciences, #4250019), CD45RO-BX017 (Akoya Biosciences, #4250023), CD163-BX069 (Akoya Biosciences, #4250079), Granzyme B-BX041 (Akoya Biosciences, #4250055), CD21-BX032 (Akoya Biosciences, #4450027), CD44-BX005 (Akoya Biosciences, #4450041), CD34-BX025 (Akoya Biosciences, #4250057), Podoplanin-BX023 (Akoya Biosciences, #4250004), PD-L1(Akoya Biosciences, #4550072), CD68 (Akoya Biosciences, #4550113), CD4 (Akoya Biosciences, #4550112), HLA-DR (Akoya Biosciences, #4550118), FOXP3 (Akoya Biosciences, #4550071), Collagen IV (Akoya Biosciences, #4550122), CD11c (Akoya Biosciences, #4550114), ICOS (Akoya Biosciences, #4550117), CD3e (Akoya Biosciences, #4550119), LAG3 (Akoya Biosciences, #4550058), CD45 (Akoya Biosciences, #4550121), PD-1 (Akoya Biosciences, #4550038), IDO1 (Akoya Biosciences, #4550123), CD20 (Akoya Biosciences, #4450018), CD31 (Akoya Biosciences, #4450017), SMA (Akoya Biosciences, #4450049), Vimentin (Akoya Biosciences, #4450050), HLA-A (Akoya Biosciences, #4450046), Pan-Cytokeratin (Akoya Biosciences, #4450020), and CD57-BX029 (BioLegend, #35602, San Diego, CA, USA) (Akoya Biosciences, #5250005).

The antibodies were kept on ice until use. The antibody cocktail stock solution was brought up to a total of 300 μL by adding the following reagents in a 1.5 mL microcentrifuge tube: staining buffer, N-blocker, G-blocker, J-blocker, and S-blocker (Akoya Biosciences, #7000017). After setting aside 200 μL of the antibody cocktail stock solution, 57 μL of the solution was removed, which is equivalent to the total volume of antibody to be added. The appropriate volume of each PhenoCycler Antibody was added to the antibody cocktail stock solution, resulting in a final solution volume of 200 μL (Table 1).

The remaining 100 μL of antibody cocktail stock solution was to prepare antibodies at a dilution ratio of 1:500. Briefly, one μL of the selected antibodies was pipetted from each vial and then diluted with the antibody cocktail stock solution to a total volume of 5 μL. Next, two μL of the diluted solution was taken and used for the final staining solution. After all the antibodies were applied, the tube was gently vortexed. 190 μL of the prepared antibody cocktail solution was drawn and quickly dispensed onto the sample slide, carefully covering the entire tissue. The sample slide was then incubated for 3 hours at room temperature on the rocker set at 30 rpm to ensure uniform staining. Following the incubation step, the sample slide was subsequently fixed with 1.6% PFA Post-Staining Fixing Solution and PhenoCycler Fixative Reagent (Akoya Biosciences, #7000017), and then washed with 1x PBS before use.

**Reporter Plate Design and Preparation**

The reporter stock solution was initially prepared in a 15 mL amber tube, considering the total number of cycles for the experiment. Thus, the reporter stock solution required for 16 cycles was designed using the following reagents: 1x buffer for

PhenoCycler with Buffer Additive (Akoya Biosciences, #700019), Assay Reagent (Akoya Biosciences, #7000002), and Nuclear Stain (Akoya Biosciences, #7000003). The total volume of the solution was 4.8 mL. After adding the reagents, the reporter stock solution was gently mixed by inverting the tube several times. Next, the reporter master mix was prepared by aliquoting the Reporter Stock Solution into separate tubes based on the number of reporters to be revealed for each corresponding cycle. Five µL of reporter for each antibody was added to the corresponding cycle master mix, resulting in a total volume of 250 µL. Once all the Reporter Master Mix had been prepared, 245 µL of the mixed solution for each cycle was transferred into the corresponding well in a 96-well plate (Akoya Biosciences, #7000006). Filled wells were then covered with a foil plate seal (Akoya Biosciences, #7000007).

**Flow Cell Assembly and PhenoCycler-Fusion**

The sample slide was washed with 1 × PBS, and the area around the tissue was cleaned to remove any excess buffer. The flow cell was attached using the Flow Cell Assembly Device (Akoya Biosciences, #240205). The flow cell-attached sample slide was then incubated in 1 × PhenoCycler Buffer + Additive for 10 minutes. Meanwhile, the PhenoCycler Designer & Fusion Software, corresponding to its experiment plan, was set up. All the required reagents and buffers for the PhenoCycler were prepared following the instructions provided by Akoya Biosciences. After incubation, the sample slide was loaded onto the PhenoCycler-Fusion for Multiplexed imaging.

**Image Preprocessing**

Raw images of the TMA slide were captured using the PhenoCycler-Fusion. We preprocessed the raw data images using Fusion software, version 1.0.6. The scan resolution was set at 0.5076 μm/pixel (20×). Saturation protection was only enabled for the DAPI setting. Using the raw and intermediate images from all 15 cycles obtained through the PhenoImager and the Fusion software, image preprocessing was performed. This process involved calculating the average autofluorescence signal of Atto550, Cy5, and AF750 in the first and last raw images ($1^{st}$ and $15^{th}$ cycles) and applying it to the final multiplexed image to remove autofluorescence. For cycle alignment, the nuclear stain pattern was compared to that of the reference cycle. The calculated offsets are used to align all cycles through rigid translation. The Fusion software subtracted the background signal from the signal channels by utilizing both the start and end blank channels. This was done to compensate for any variations in autofluorescence during a PhenoCycler experiment. As a result, we generated a multichannel Qptiff image for a single TMA slide that is devoid of autofluorescence and contains all image layers and metadata. We converted the Qptiff file into an ome.tiff image file using the open-source software QuPath[19] (Figure S2B).

**Dimensionality Reduction**

Using cuML (https://github.com/rapidsai/cuml), the RAPIDS equivalent of scikit-learn, we performed dimensionality reduction by calculating the t-SNE (t-Distributed Stochastic Neighbor Embedding) coordinates. In this process, we set the perplexity parameter to 60 and determined N_Neighbors to be three times the perplexity value. Subsequently, we created t-SNE plots to visualize the results of the dimensionality reduction analysis.

### ROI Area Calculation

To calculate the area of each Region of Interest (ROI), we utilized a combination of image processing techniques using the DAPI, Vimentin, αSMA (α-smooth muscle actin), and Collagen IV channels. The process involved the following steps: (1) image combination, in which we combined the aforementioned channels to create a composite image; (2) thresholding, in which Otsu's thresholding method from the skimage.filters (https://scikit-image.org/) was applied to segment the image into foreground and background; (3) binary dilation, in which we used binary dilation with skimage.morphology.binary_dilation to enhance the regions of interest; (4) filling holes, in which any holes in the binary image were filled using skimage.morphology.binary_fill_holes; and (5) removing small objects, in which small objects in the binary image were removed using the skimage.morphology.remove_small_objects function. After performing these operations, we obtained a binary image with clearly defined areas of interest (Figure S5). We used NumPy (https://numpy.org) to aggregate these areas and ultimately computed the total area.

### Proportion and Density Calculation

We calculated the proportion by dividing the number of cells within a specific cluster by the total cell count. Density was determined by dividing the cell count of the respective cluster by the area size of the corresponding ROI.

**Table S1**. Dilution Ratio and Volume Requirement of Antibodies for Staining Solution

| Antibody | Barcode | Dilution Ratio | Antibody | Barcode | Dilution Ratio | Antibody | Barcode | Dilution Ratio |
|---|---|---|---|---|---|---|---|---|
| CD8 | BX026 | 1:100 | PD-L1 | BX043 | 1:200 | CD20 | BX007 | 1:200 |
| E-cadherin | BX014 | 1:200 | CD68 | BX015 | 1:500 | CD31 | BX001 | 1:100 |
| CD14 | BX037 | 1:500 | CD4 | BX003 | 1:100 | SMA | BX013 | 1:500 |
| Ki67 | BX047 | 1:500 | HLA-DR | BX033 | 1:200 | Vimentin | BX022 | 1:500 |
| CD45RO | BX017 | 1:200 | FOXP3 | BX031 | 1:200 | HLA-A | BX004 | 1:200 |
| CD163 | BX069 | 1:500 | Collagen IV | BX042 | 1:200 | PanCK | BX019 | 1:500 |
| Granzyme B | BX041 | 1:200 | CD11c | BX024 | 1:500 | IDO1 | BX027 | 1:500 |
| CD21 | BX032 | 1:500 | ICOS | BX054 | 1:500 | CD57 | BX029 | 1:50 |
| CD44 | BX005 | 1:200 | CD3e | BX045 | 1:200 | PD-1 | BX046 | 1:500 |
| CD34 | BX025 | 1:500 | LAG3 | BX055 | 1:500 | | | |
| Podoplanin | BX023 | 1:500 | CD45 | BX021 | 1:500 | | | |

**Table S2**. Patient characteristics

| Characteristics | High-CRP (patient n = 6) | Low-CRP (patient n = 3) | Normal aorta (patient n = 3) |
|---|---|---|---|
| Age, y | 65.5 ± 6.4 | 62.7 ± 3.2 | 37.3 ± 23.7 |
| Male, n (%) | 6 (100.0) | 3 (100.0) | 3 (100.0) |
| Body mass index, kg/m2 | 24.0 ± 3.8 | 24.6 ± 4.8 | 16.0 ± 0.6 |
| Serum CRP level, mg/dL | 0.9 ± 0.8 | 0.1 ± 0.0 | 2.2 ± 2.9 |
| Abdominal aorta size, cm | 6.3 ± 1.5 | 5.8 ± 0.3 | NA |
| Diabetes mellitus, n (%) | 1 (16.7) | 1 (33.3) | 1 (33.3) |
| Hypertension, n (%) | 4 (66.7) | 3 (100.0) | 1 (33.3) |
| Previous Cardiovascular diseasea, n (%) | 2 (33.3) | 1 (33.3) | 3 (100.0) |
| Alcohol, n (%) | 2 (33.3) | 3 (100.0) | 0 (0.0) |
| Smoking, n (%) | 6 (100.0) | 2 (66.6) | 0 (0.0) |
| Dyslipidemiab, n (%) | 6 (100.0) | 2 (66.6) | 0 (0.0) |
| Chronic kidney disease, n (%) | 0 (0.0) | 0 (0.0) | 1 (33.3) |
| Arteriosclerosis obliterans, n (%) | 0 (0.0) | 2 (66.6) | 0 (0.0) |
| Preoperative Diagnosis, n (%) | | | |
| Abdominal aortic aneurysm | 7 (100.0) | 3 (100.0) | 0 (0.0) |
| Dilated cardiomyopathy | 0 (0.0) | 0 (0.0) | 2 (66.6) |
| Hypertrophic cardiomyopathy | 0 (0.0) | 0 (0.0) | 1 (33.3) |

*Included history of stroke, myocardial infarction, or angina requiring percutaneous coronary intervention

†Patients who were on statin medication

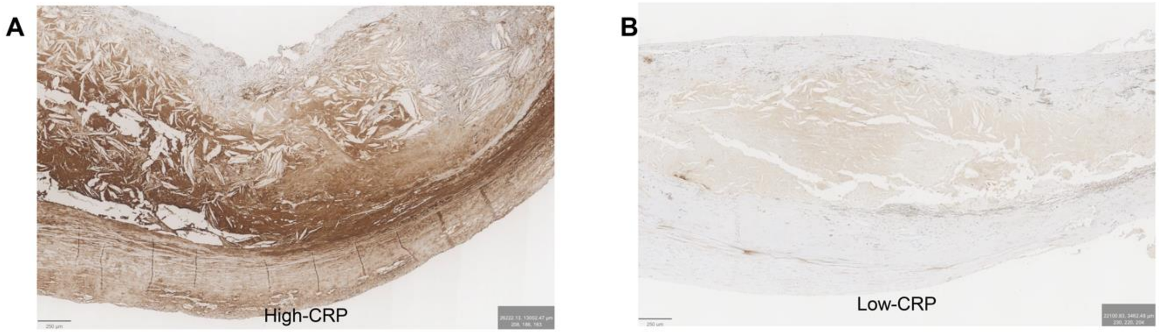


**Figure S1**. Representative anti-CRP antibody immunohistochemical staining of the High-CRP (left) and Low-CRP (right). This anti-CRP antibody detects both pentameric CRP and monomeric CRP. The High-CRP group showed diffuse and strong immunopositivity for the anti-CRP antibody, while the Low-CRP group showed negativity for the anti-CRP antibody.

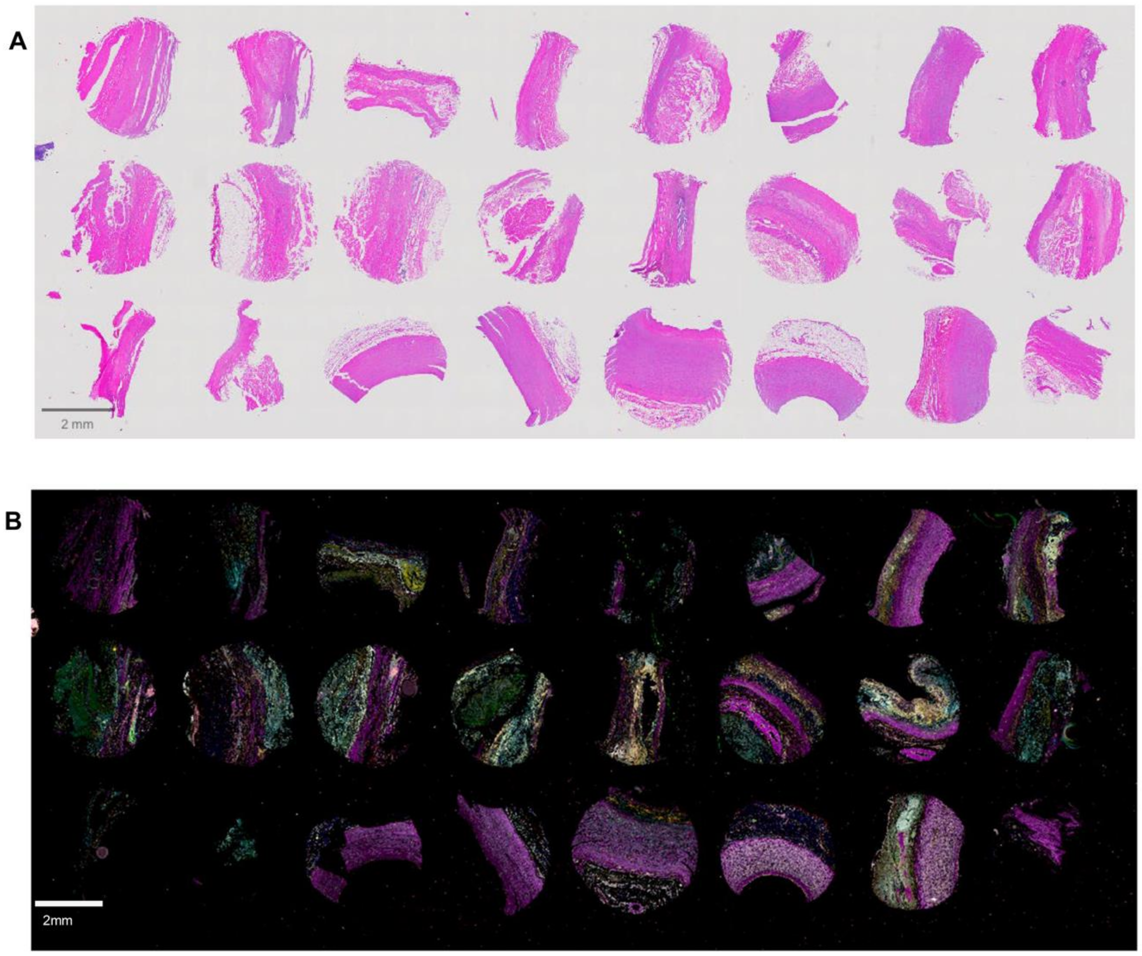


**Figure S2. Scan image of Tissue microarray (TMA)**

(A) Digitalized image of hematoxylin and eosin staining

(B) Scan view of CODEX multiplexed imaging

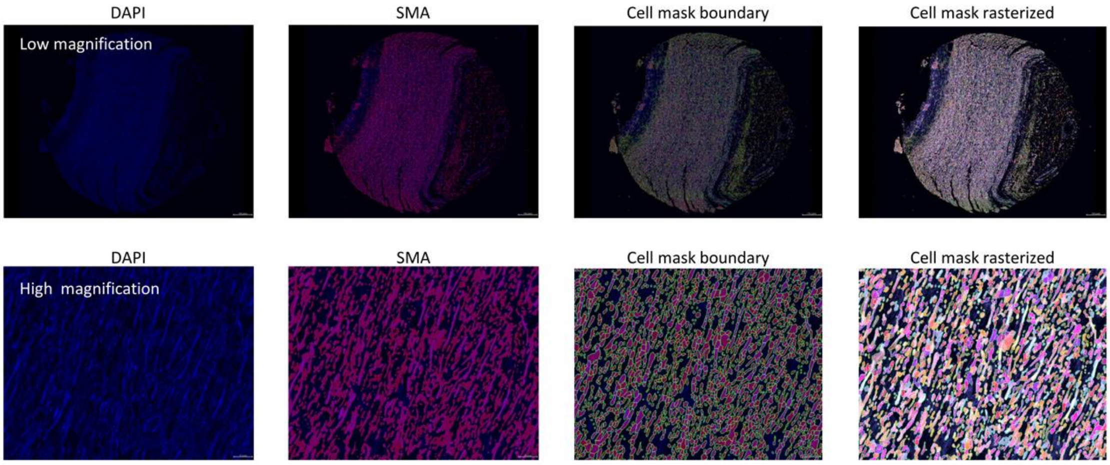


**Figure S3**. Cell segmentation mask visualization on normal aortic wall.

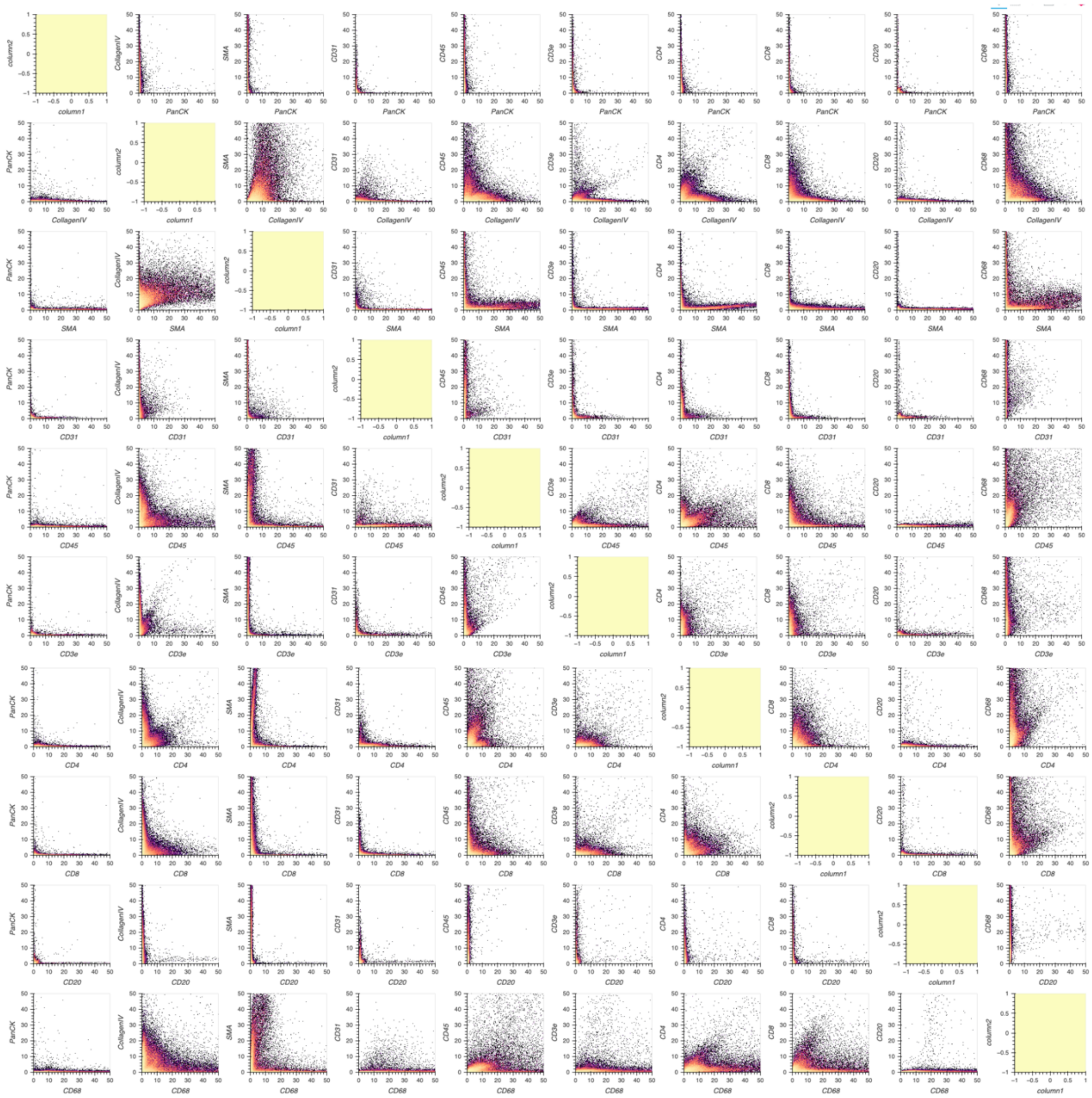

**Figure S4**. Signal intensity scatter plot.

We performed scatter plots for antibodies known to exhibit mutually exclusive expression patterns. This set of antibodies formed an L-shaped plot. Additionally, epithelial markers were negative in vascular samples. These findings indicate the proper staining of the antibodies.

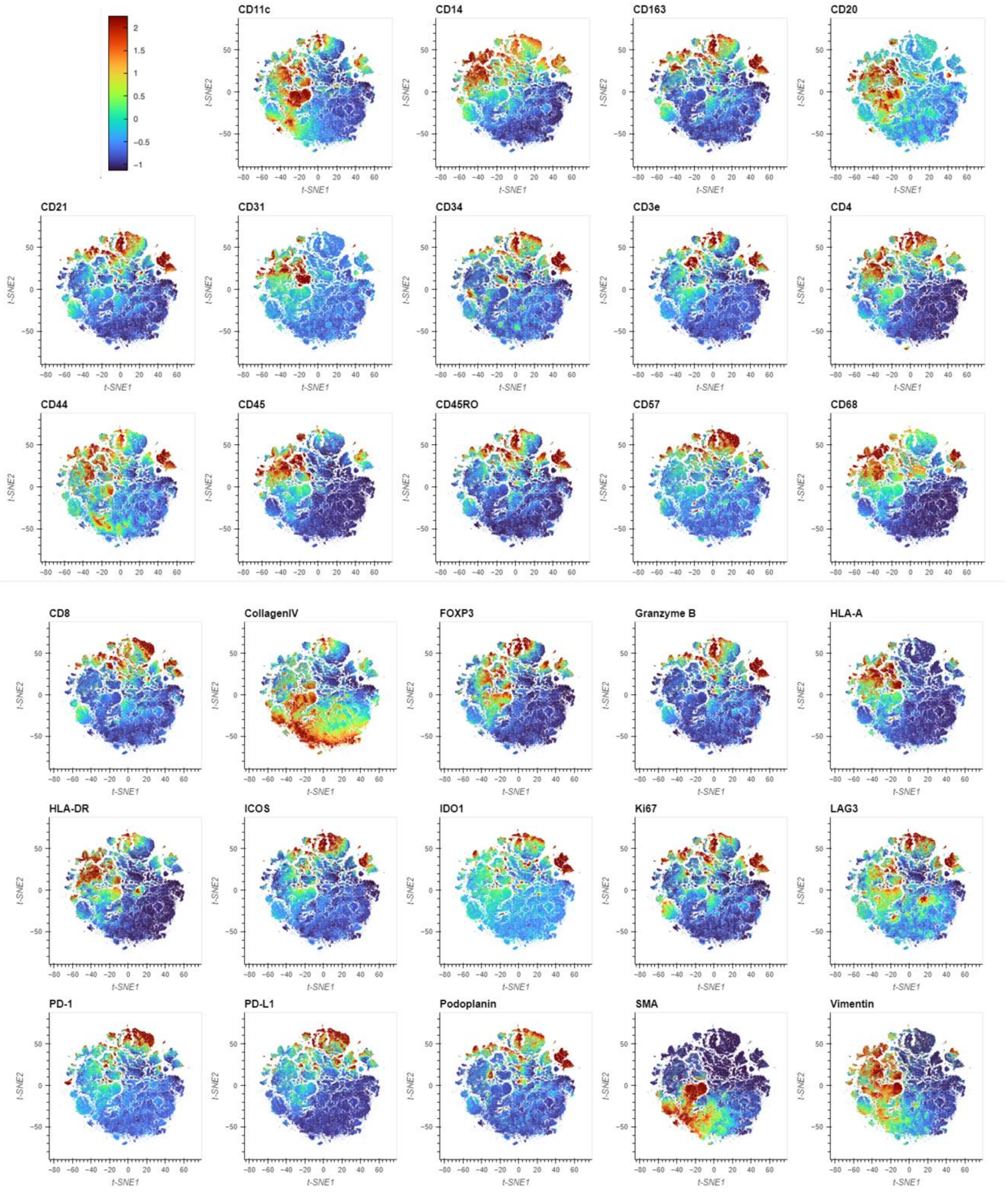


**Figure S5**. In the t-SNE plot, the normalized expression of each marker was overlaid.

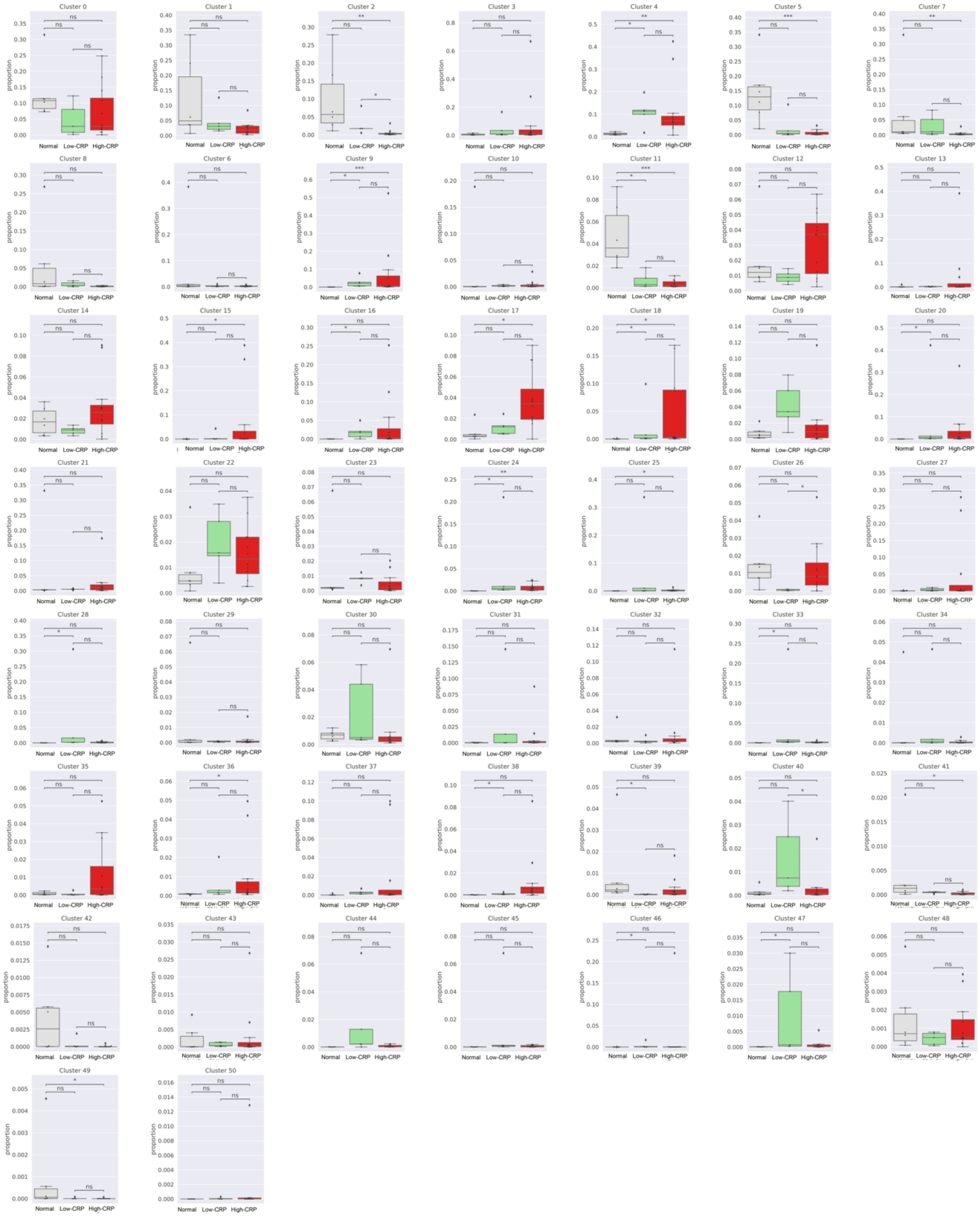

**Figure S6**. Cell proportion comparison. Twenty-four specific cell clusters (clusters 2, 4, 5, 7, 9, 11, 15, 16, 17, 18, 20, 24, 25, 26, 28, 33, 36, 38, 39, 40, 41, 46, 47, 49) were significantly more frequently observed in the AAA-high or Low-CRP groups.

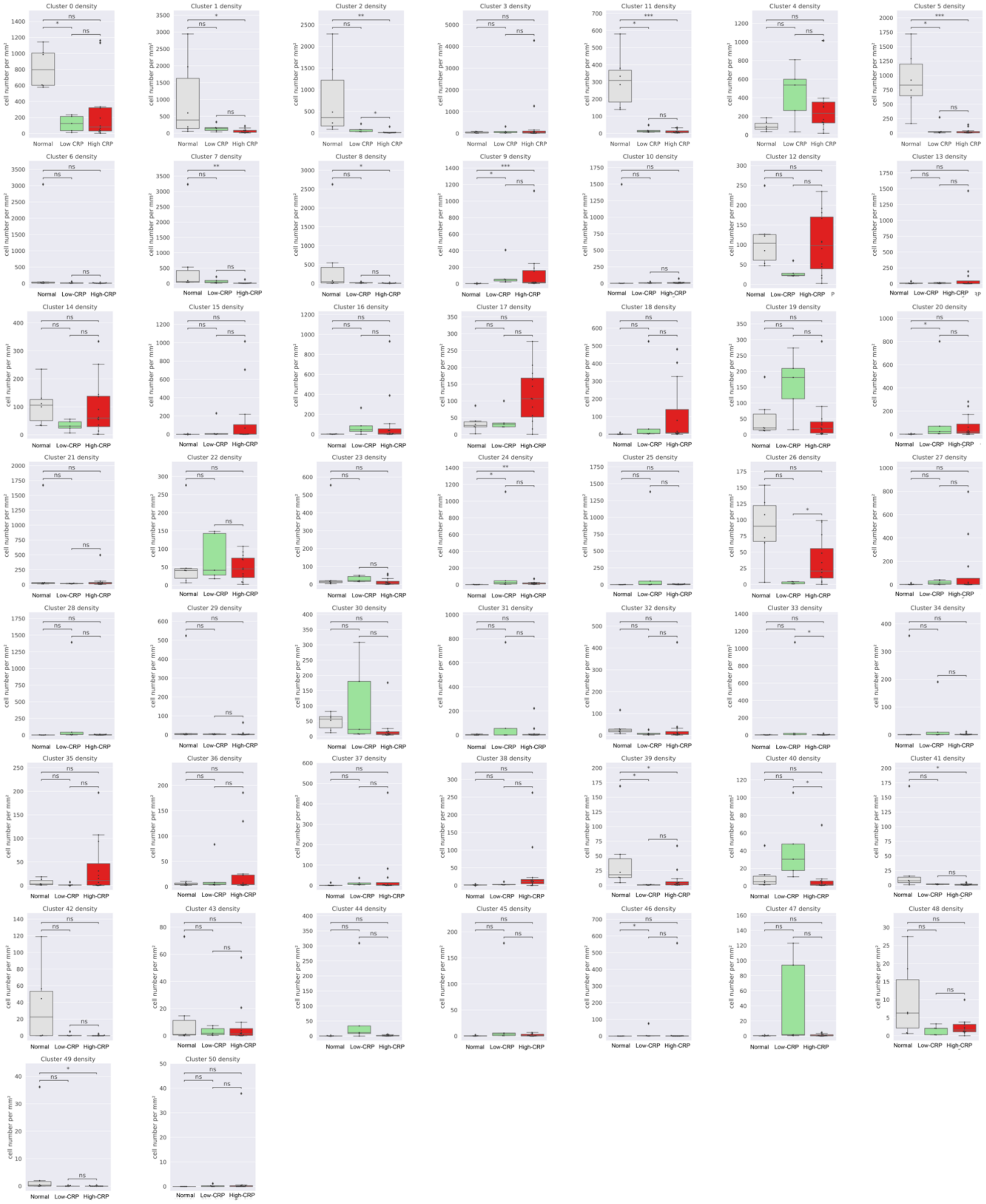


**Figure S7**. Comparison of cell density [cell number/mm$^2$, Immune or stromal cell number/area of each ROI] in normal aorta, Low-CRP, and High-CRP groups. The trend was similar to the composition comparison graph in Figure S6.

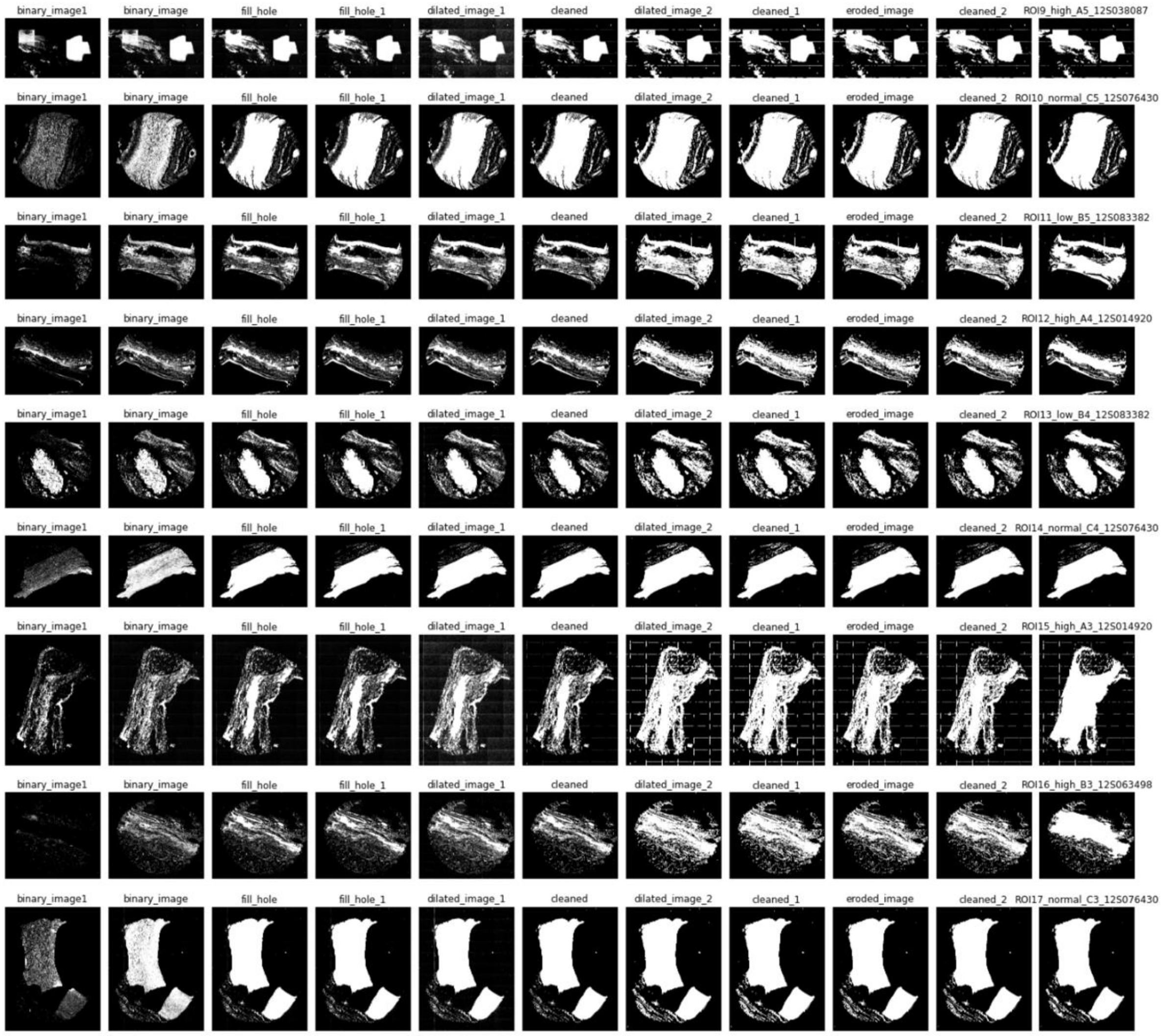


**Figure S8**. Creation of binary images and calculation of area through SMA image modification.
